# Germaphobia! Does our Relationship with, and Knowledge of Biodiversity Affect our Attitudes Towards Microbes?

**DOI:** 10.1101/2021.02.08.430200

**Authors:** Jake M. Robinson, Ross Cameron, Anna Jorgensen

## Abstract

Germaphobia –– a pathological aversion to microorganisms –– could be contributing to an explosion in human immune-related disorders via mass sterilisation of surfaces and reduced exposure to biodiversity. Loss of biodiversity and our connectedness to nature, along with poor microbial literacy may be augmenting the negative consequences of germaphobia on ecosystem health. In this study, we created an online questionnaire to acquire data on attitudes towards, and knowledge of microbes. We collected data on nature connectedness and interactions with nature and explored the relationships between these variables. We found a significant association between attitudes towards microbes and both duration and frequency of visits to natural environments. A higher frequency of visits to nature per week, and a longer duration spent in nature per visit, significantly associated with positive attitudes towards microbes. We found no association between nature connectedness and attitudes towards microbes. We found a significant relationship between knowledge of ‘lesser known’ microbial groups (e.g., identifying that fungi, algae, protozoa, and archaea are microbes) and positive attitudes towards microbes. However, we also found that people who correctly identified viruses as being microbes expressed less positive views of microbes overall –– this could potentially be attributed to a ‘COVID-19 effect’. Our results suggest that basic microbial literacy and nature engagement may be important in reducing/preventing germaphobia. The results also suggest that a virus-centric phenomenon (e.g., COVID-19) could increase broader germaphobia. As the rise of immune-related disorders and mental health conditions have been linked to germaphobia, reduced biodiversity, and non-targeted sterilisation, our findings point to a feasible strategy to potentially help ameliorate these negative consequences. A greater emphasis on microbial literacy and promoting time spent in nature could be useful in promoting resilience in human health and more positive/constructive attitudes towards the foundations of our ecosystems – the microorganisms.

## 1. Introduction

Germaphobia – also known as ‘mysophobia’ – is the pathological fear of, and aversion to dirt and microorganisms (henceforth referred to as ‘microbes’) (Zemke et al. 2015). The rise of germaphobia has likely been influenced by decades of advertising campaigns creating negative perceptions of microbes, and falsely prompting mass (non-targeted) sterilisation of surfaces to achieve ‘safe’ human environments (Timmis et al. 2019). Symptoms of germaphobia include excessively washing hands, over-use of sanitisers and antibiotics and avoiding certain places due to perceived to fear of microbial exposure (Qadir and Yameen, 2019). However, far less than 1% of the microbes on the planet are human pathogens (Zobell and Rittenberg, 2011; Balloux and van Dorp, 2017). Moreover, germaphobia may have contributed to the current explosion in human immune-related disorders (such as diabetes, asthma, and inflammatory bowel disease) (Jun et al. 2018; Timmis et al. 2019). This is thought to be attributed to the notion that exposure to environmental microbiomes – the diverse network of microbes in a given environment – plays an important role in human health (Rook et al. 2003; Dannemiller et al. 2014; Stein et al. 2016; Arleevskaya et al. 2019; Liddicoat et al. 2019; Selway et al. 2020). Indeed, from a young age, exposure to a diverse range of environmental microbes is considered to be essential for the training and regulation of our immune systems (Flies et al. 2020; Renz and Skevaki, 2020; Roslund et al. 2020). A stable and functional human microbiome is colonised following birth. Firstly by the mother’s skin and breast milk, and later supplemented from visitors, pets, biodiverse environments, and a ‘normal dirty’ (not overly cleaned) home environment (DeWeerdt, 2018). Germaphobia could conceivably inhibit all of these activities (e.g., avoiding playing in soil or staying away from animals), and if the microbiome assembly process is derailed, the health consequences could be long-term.

Against the backdrop of COVID-19 –– a situation that could conceivably increase germaphobia –– in addition to being hygienic, we need to promote the concept that the majority of microbes are in fact innocuous and/or beneficial to human health.

Microbial communities and their interactions also play essential roles in carbon and nutrient cycling, climate regulation, animal and plant health, and global food security (Cavicchioli et al. 2019; Li et al. 2020; Trivedi et al. 2020). Therefore, microbial biodiversity is of vital importance for the ability of ecosystems to simultaneously provide multiple ecosystem services (Guerra et al. 2020). Consequently, ongoing degradation of microbial communities poses an existential threat to global macro-level biodiversity and to human societies across the planet. Loss of biodiversity and our affective, cognitive and experiential connection with the natural world (also known as ‘nature connectedness’), along with poor microbial literacy (such as awareness of the different types of microbes and their importance) may be augmenting the negative consequences of germaphobia on ecosystem health (Cavicchioli et al. 2019; Robinson and Breed, 2020). Is our diminishing connection with (the rest of) the natural world helping to drive germaphobia itself? This could have considerable implications for the health of humans and that of our diverse ecosystems.

In this study, we used an online questionnaire to acquire data on attitudes towards microbes. We collected data on nature connectedness using the Nature Relatedness 6 Scale – a validated psychological instrument (Nisbet et al. 2013), and data on respondents’ interactions with nature (including typical duration and frequency of visits to nature). To gauge respondents’ basic knowledge of microbes, we asked them to select all of the organisms (from a list) that they considered to be microbes. The relationships between these variables were then assessed using a range of statistical methods including logistic regression models, Mann Whitney U tests, and 2-sample tests for equality of proportions with continuity correction in R.

The primary objectives of this study were to: **(a)** assess whether people’s patterns of exposure to nature associated with their attitudes towards microbes (i.e., a positive or negative view); **(b)** assess whether people’s level of subjective connectedness to nature associated with their attitudes towards microbes; and, **(c)** investigate whether basic knowledge of microbial groups (e.g., identifying that fungi, algae, protozoa, and archaea are also microbes) associated with attitudes towards microbes.

Gaining a better understanding of the factors that may aid in reducing/preventing germaphobia could help to inform environmental and public health policy. For example, improving microbial literacy and promoting campaigns that seek to reconnect humans with the wider biotic community could bring immense value to both human and environmental health. Microbes are the foundations of our ecosystems and are essential to the survival of all life on Earth (Cavicchioli et al. 2019). While targeted hygiene approaches and continued efforts to control infectious diseases are undoubtedly vital, germaphobia only serves to inhibit a more nuanced awareness of, and mutually-advantageous relationship with these diverse, underappreciated, and indispensable lifeforms.

## 2. Materials and Methods

### Online questionnaire

We produced a research questionnaire using the Smart Survey online software (Smart Survey, 2020). The questionnaire included 21 multi-format questions (Supplementary Materials, Appendix I). The questions were devised to gather data on respondents’ attitudes towards microbes, their nature connectedness, and interactions with nature. The online survey was active between April and July 2020. We asked participants to answer questions regarding how emotionally and cognitively connected they felt to nature using the Nature Relatedness Scale (NR-6) (Nisbet et al. 2013; Kettner et al. 2019). The NR-6 comprises 6 questions, and answers are recorded using a 1-5 Likert scale. Examples of questions include “My relationship to nature is an important part of who I am”, “My ideal vacation spot would be a remote, wilderness area”, and “I feel very connected to all living things and the earth”. Items were averaged, and higher scores indicated stronger subjective connectedness to nature. This validated instrument has been used in several previous environmental psychology studies (Nisbet et al. 2013; Obery and Bangert, 2017; Whitburn et al. 2020). We also asked several pilot-tested questions regarding typical exposure to nature such as duration and frequency of visits to natural environments. For this study ‘natural environments’ and/or ‘nature’ were considered to be less anthropogenic/built-up environments, typically containing a large proportion of vegetation and wildlife such as woodlands, parks, and meadows.

To acquire data on respondents’ attitudes towards microbes, we devised questions such as “do you consider microbes to be good?; bad?; some are good, some are bad?; or, neither are good or bad?”. We also devised a pilot-tested word-association measure using three categories: positive association, neutral association, and negative association. To reduce potential bias, the categories were not revealed to the respondents and each category contained five randomly-ordered words, displayed as one amalgamated list. In the positive category, respondents could choose from words such as ‘essential’ and/or ‘beneficial’. In the neutral category respondents could choose from words such as ‘nature’ and/or ‘mobile’. In the negative category respondents could choose from words such as ‘disease’ and/or ‘nuisance’. Respondents were asked to select a total of three words that best reflected their view of microbes. To gauge respondents’ knowledge of microbes, we asked them to select all of the organisms that they considered to be microbes. The list included bacteria, viruses, fungi, algae, protozoa, and archaea.

We also acquired key demographic information including postal code, relative deprivation, age, gender, highest level of education, and occupation. The questionnaire, along with a detailed participant information sheet and consent form was distributed across the world via a secure weblink. We used several non-random sampling methods to reach respondents including: social media posting, emailing volunteer groups, and carrying out an online search of publicly available community group directories. The only exclusion criterion for the study was: people under 18 years of age. The questionnaire was ethically reviewed by the internal review committee in the Department of Landscape Architecture at the University of Sheffield (the authors’ academic institution).

### Statistical analysis

To assess relationships between duration and frequency of visits to nature and attitudes towards microbes, we acquired a score from the word-association output by summing the positive, neutral and negative values given by each respondent. We then assigned the positive and negative scores into two groups and compared the mean duration and frequency of visits to nature of each group using the two-sample Mann-Whitney U test with continuity correction in R. We assessed proportional differences between groups, in which respondents either did or did not identify different microbial groups (i.e., bacteria, viruses, fungi, algae, protozoa, and archaea) and their respective word-association scores using the 2-sample tests for equality of proportions with continuity correction in R. We built logistic regression models to assess the relationships between nature connectedness and attitudes towards microbes. For these models, an odds ratio (OR) of 1 or above equated to the predictor variable (nature connectedness score) increasing the odds of a positive attitude towards microbes. An OR <1 equated to the predictor variable decreasing the odds of a positive attitude towards microbes. We adjusted for several covariates including age, gender, deprivation, and level of education.

## 3. Results

A total of *n* = 1184 respondents completed the questionnaire. A broad distribution of responses from across the world was acquired (Fig. 1, A); however, the main cluster (*n* = 993) was from England, UK (Fig. 1, B).

**Fig. 1.**
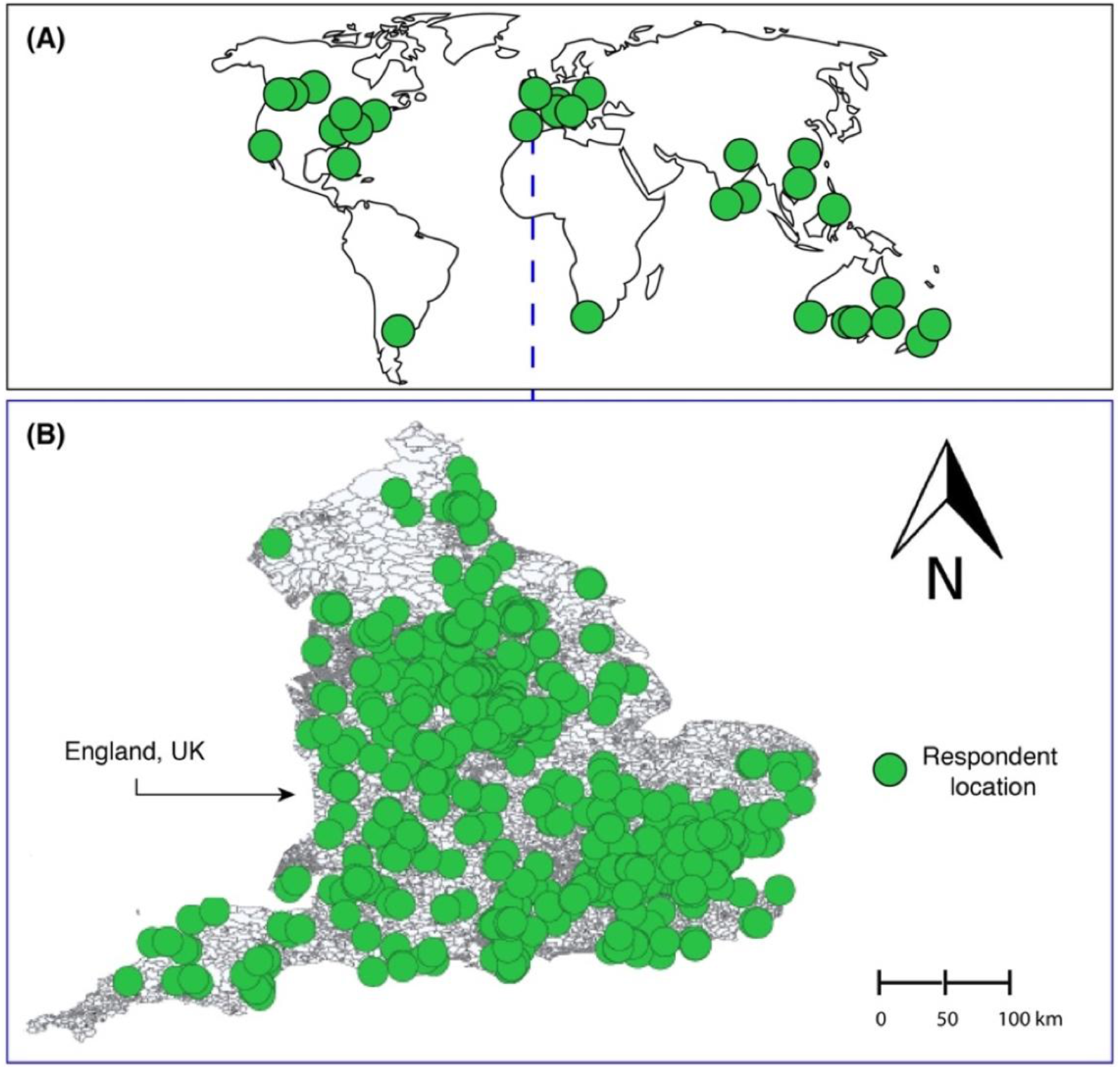
Distribution of respondents, whereby (A) shows the global distribution, and (B) shows England, UK – the geographical source of the majority of responses (*n* = 993).

Respondents who identified as being female (*n* = 851 or 72%) outnumbered those who identified as being male (*n* = 331 or 28%), trans woman (*n* = 1 or 0.1%), and non-binary (*n* = 1 or 0.1%). There was also a skew towards respondents with a higher level of education (*n* = 847 or 72% with ≥ undergraduate degree). In terms of age, the distribution either side of the median was similar (*n* = 624 or 53% were ≥55 years old; and *n* = 560 or 47% were ≤54 years old).

### Duration in and frequency of visits to natural environments, and attitudes towards microbes

Our results show that respondents with a net positive word-association score for microbes (i.e., those who viewed microbes more positively) spent significantly more time per visit (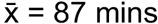) to natural environments such as woodlands, parks, and meadows compared to respondents with a net negative word-association score for microbes (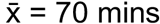) (W = 3995, *p* = <0.01) (Fig. 2).

**Fig. 2.**
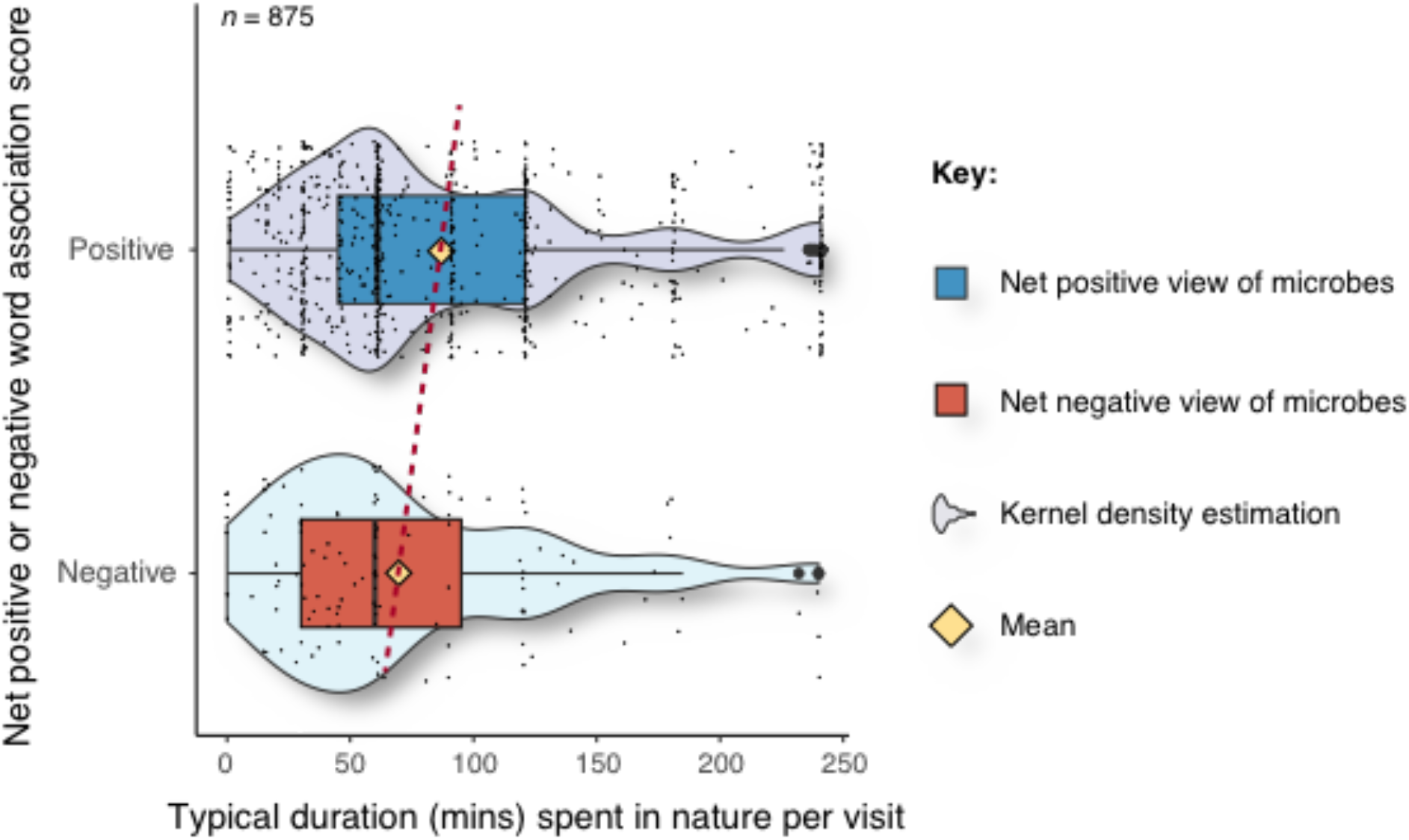
Typical duration spent in natural environments per visit for respondents with net positive and net negative word-association scores. The yellow diamond represents the mean value.

Our results also show that respondents with a net positive word-association score for microbes visited natural environments such as woodlands, parks, and meadows significantly more often (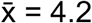 visits in a given week) compared to respondents with a net negative word-association score for microbes (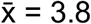 visits in a given week) (W = 3935, *p* = <0.01) (Fig. 3).

**Fig. 3.**
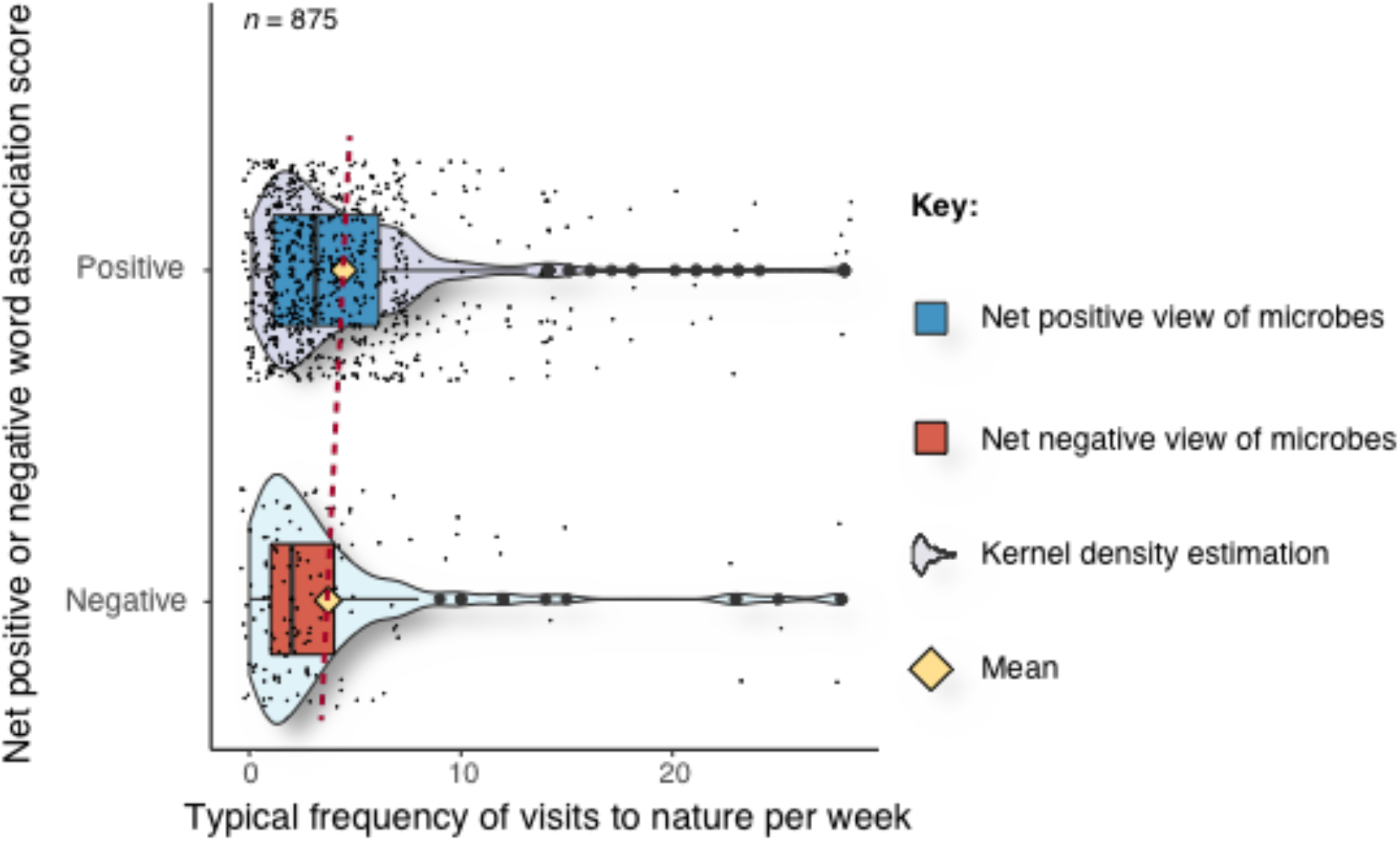
Typical frequency of visits to natural environments per week for respondents with net positive and net negative word-association scores. The yellow diamond represents the mean value.

**Fig. 3.**
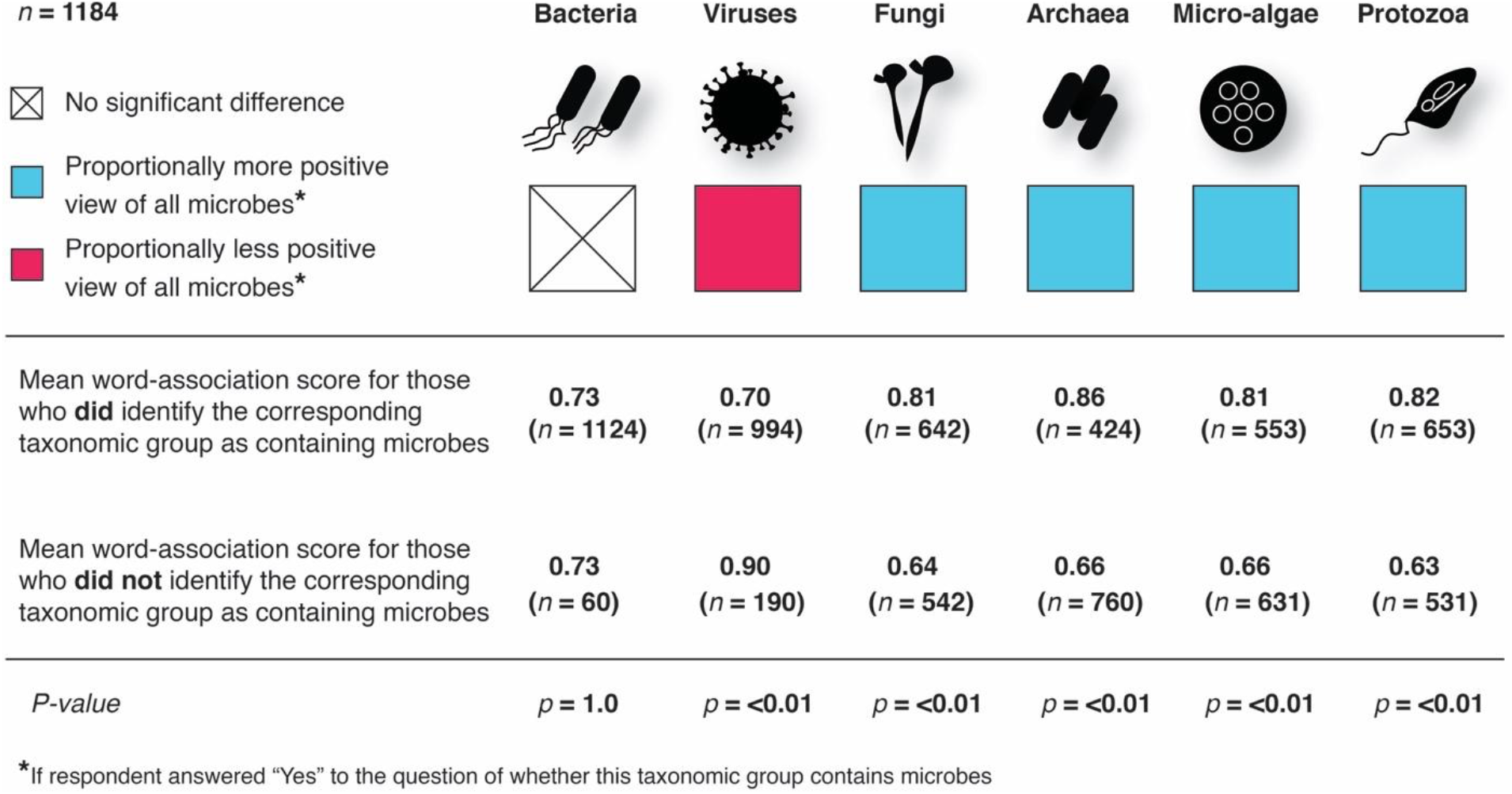
Differences in mean microbe word-associated scores for respondents who correctly identified a given taxa as being a microbe compared to those who did not identify the taxa as being a microbe. There were significantly higher (in positivity) word-association scores (indicated by the blue boxes) for respondents who correctly identified that fungi, archaea, micro-algae, and protozoa are microbes compared to those who did not.

### Nature connectedness and attitudes towards microbes

We found no association between nature connectedness (measured using the NR-6 Scale) and attitudes towards viruses (OR: 0.99 (0.95, 1.02) *p* = 0.54) or all other microbes (OR: 1.01 (0.89, 1.16) *p* = 0.86) (Table 1).

**Table 1.**
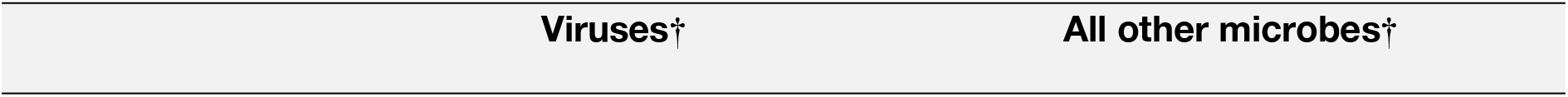
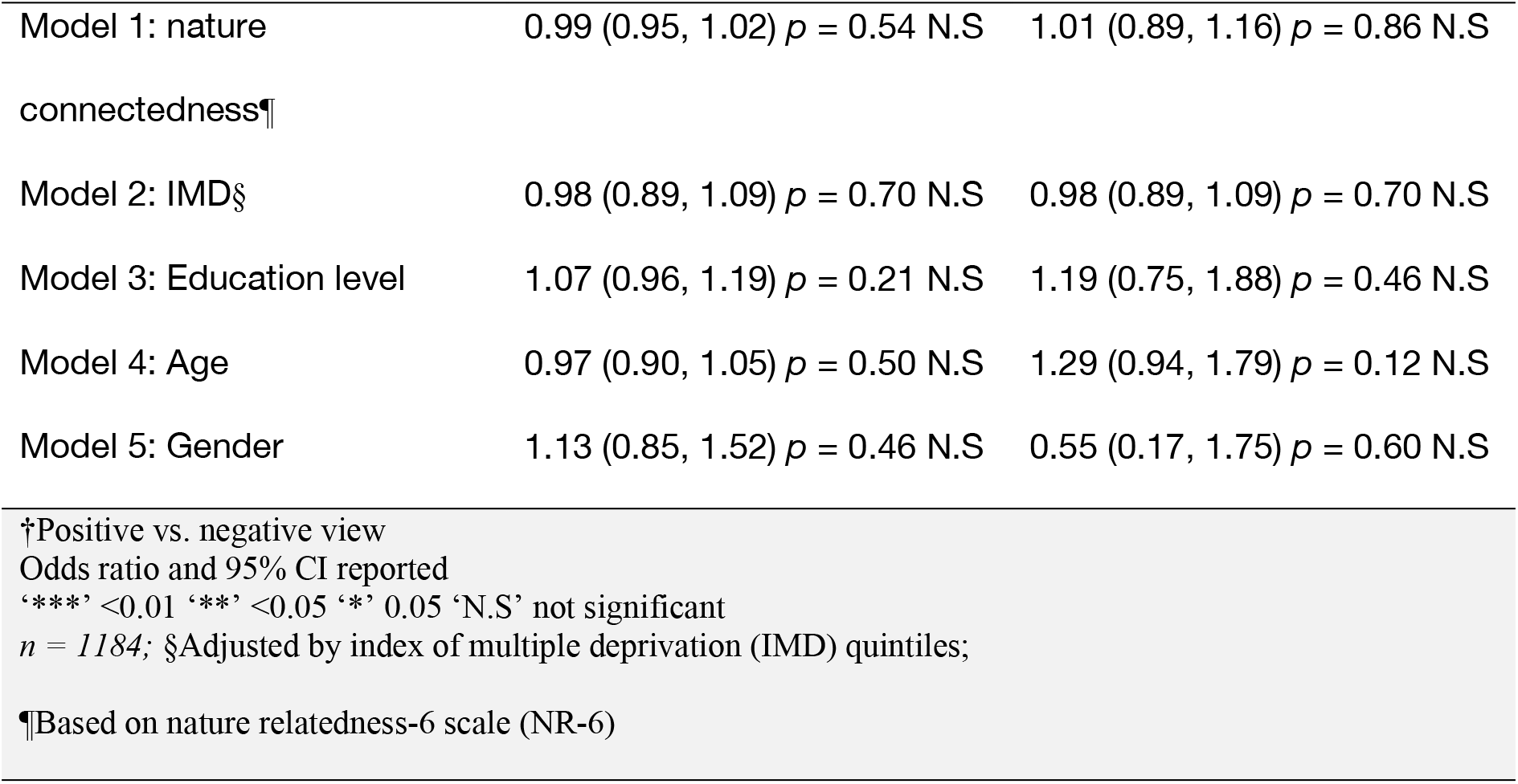
Associations between positive or negative views of microbes and nature connectedness, adjusting for relative deprivation, education, age and gender.

### Microbial literacy and attitudes towards microbes

Mean positive scores (derived from word-association) towards all microbes were significantly higher for those who correctly identified that fungi (*X^2^* = 42.5, df = 1, *p* = <0.01) archaea (*X^2^* = 52, df = 1, *p* = <0.01) micro-algae (*X^2^* = 30, df = 1, *p* = <0.01) and protozoa (*X^2^* = 51, df = 1, *p* = <0.01) were microbes compared to those who did not identify these groups as being microbes. Mean positive scores towards all microbes were significantly lower for those who correctly identified that viruses were microbes compared to those who did not identify viruses as being microbes (*X^2^* = 30.7, df = 1, *p* = <0.01). There were no significant differences in scores between respondents who correctly identified bacteria as being microbes (*n* = 1124) compared to those who did not (*n* = 60) (*X^2^* = <0.01, df = 1, *p* = 1.0).

## 4. Discussion

This study shows a significant relationship between our attitudes towards microbes, how long we spend in natural environments and how often we visit them. However, we found no association between nature connectedness (one’s affective, cognitive and experiential connection with the natural world) (Cheung et al. 2020; Choe et al. 2020) and attitudes towards microbes. Importantly, we found a significant relationship between knowledge of ‘lesser known’ microbial groups (e.g., identifying that fungi, algae, protozoa, and archaea are microbes) and positive attitudes towards microbes. This study suggests that basic microbial literacy and nature exposure may be important in reducing/preventing germaphobia.

As mentioned, a higher frequency of visits to nature per week, and a longer duration spent in nature per visit, significantly associated with positive attitudes towards microbes. It is important to note that the directionality of the relationship is unknown (i.e., whether spending more time in nature helps to establish more positive attitudes towards microbes, or whether other factors related to more positive attitudes increase the likelihood of spending more time in nature). Conceivably, being less averse to microbes could increase one’s desire to spend time in environments with natural features such as plants and soil – key sources of dense microbial communities (Liddicoat et al. 2019; Robinson et al. 2020). On the other hand, a greater habituation to these kinds of environments and an affinity for diverse life-forms could conceivably reduce one’s aversion to microbes in general. It is important to acknowledge here that spending time in natural environments exposes us to a diverse suite of microbial communities (Robinson et al. 2020; Selway et al. 2020) that are thought to have important beneficial effects on our health (Haahtela, 2019; Renz and Skevaki, 2020). Therefore, whatever the actual directionality of the proposed relationship is, it is likely to have an important impact on our health and could help to ameliorate the negative consequences of germaphobia. In one direction (i.e., contingent on factors related to more positive attitudes towards microbes increasing the likelihood that we will spend more time in nature), we could potentially gain the many benefits associated with nature engagement. These include improvements in immune health (Li et al. 2010; Rook, 2013), mental health (Birch et al. 2020; Callaghan et al. 2020), and cardiovascular health (Yao et al. 2020; Yeager et al. 2020). In the alternative direction (i.e., spending more time in natural environments which may help to establish more positive attitudes towards microbes), our positive attitudes towards microbes could conceivably reduce the likelihood that we carry out mass (non-targeted) sterilisation of our local environments, which could also have important implications for our health (Jun et al. 2018; Parks et al. 2020; Prescott, 2020; Renz and Skevaki, 2020). This relationship could also be non-dichotomous (or potentially even a virtuous loop) in the sense that our positive attitudes towards microbes may predispose us to spend more time in nature––an act that may enhance our positive attitudes towards microbes, and the feedback continues. This theoretical relationship warrants further research.

There is potentially an important systematic error to also consider here––recently termed the ‘*Holobiont Blindspot’* (Robinson and Cameron, 2020). The microbes within our bodies could influence our decisions to spend time in particular environments and/or select conditions that favour particular taxa within the human microbiome, which manifests as a cognitive bias if unrecognised. Could changes in our microbiome influence our germaphobia? Further research is needed.

Given that nature engagement associates with positive attitudes towards microbes, it would perhaps be expected that nature connectedness may also associate with positive attitudes towards microbes. Studies have shown that people who exhibit higher levels of nature connectedness are more likely to spend time in and engage with natural environments (Capaldi et al. 2014; Capaldi et al. 2015), and reciprocally, spending time in nature can enhance one’s nature connectedness (Nisbet et al. 2019; Chawla, 2020). However, the results of our study show that no significant relationship existed between the nature connectedness of our respondents and their attitudes towards microbes. This could be confounded by other factors, however, age, gender, education and deprivation were controlled for with similar non- significnt results. It may simply be that one’s affective, cognitive and experiential connection with nature is not an important factor in predicting one’s attitude towards microbes. We can only speculate and say that the invisibility of microbes to the human eye could conceivably negate the affective, cognitive and experiential connection that one may establish with, for example, charismatic fauna or aesthetically-appealing flora. Alternatively, this result could be a facet of the nature connectedness instrument used (the NR-6 Scale). Perhaps a more detailed version of the instrument such as the 17-item Connectedness to Nature Scale (CNS) (Mayer and Frantz, 2004) would reveal alternative findings. This warrants further research.

Finally, our study shows a significant relationship between basic level of microbial literacy and attitudes towards microbes. Respondents who correctly identified that lesser publicised (as microbes) organisms –– such as algae, fungi, archaea, and protozoa –– were microbes, showed higher positivity scores towards microbes. This implies that basic microbial literacy may be an important factor in the formation of one’s attitudes towards microbes, and thus could influence the onset of germaphobia. Interestingly, mean positive scores towards all microbes were significantly lower for those who correctly identified that viruses were microbes compared to those who did not identify viruses as being microbes. Although further research is needed, one explanation could be that the COVID-19 (virus) pandemic had an effect on people’s overall view of microbes. This is unsurprising given the damage the pandemic has cause and the multi-pronged approach taken to try and eliminate the SARS-CoV-2 virus. However, it could conceivably have negative cascading effects on our health by contributing to broader germaphobia.

Microbes are the foundations of our ecosystems and are essential to the survival of all life on Earth (Cavicchioli et al. 2019). We now have the technology to easily characterise and learn about these diverse invisible communities that continuously surround us, providing essential ecosystem services. Perhaps in an educational context, greater emphasis can be placed on microbial literacy moving into the future. With a more nuanced awareness of, and mutually-advantageous relationship with these diverse, underappreciated, and indispensable lifeforms, germaphobia can potentially be reduced, while still maintaining the critically important targeted-hygiene and efforts to control infectious diseases.

### Limitations

Our study has some important limitations. Firstly, the results in the study are correlational. Therefore, strict inferences of causation are not possible. Along similar lines, inferences regarding the directionality of the relationships are also not possible. Non-random sampling methods were used in this study. This means accurate calculations of error and representativeness are not possible. Perhaps one of the most important limitations is that self-reported data collection methods come with inherent biases. For example, responder bias –– where participants (either intentionally or by accident) choose an untruthful or inaccurate answer. Further controlled research is required to fully unravel the complexities of the observed relationships.

## 5. Conclusions

This study suggests that basic microbial literacy and nature exposure may be important in reducing/preventing germaphobia. As the rise of immune-related disorders and mental health conditions have been linked to germaphobia, reduced biodiversity, and non-targeted sterilisation, our findings point to a simple strategy to potentially help ameliorate these negative consequences. Indeed, a greater emphasis on microbial literacy and promoting time spent in nature could be useful in promoting resilience in human health and more positive/constructive attitudes towards the foundations of our ecosystems – the microorganisms.

## Author Contributions

J.M.R, R.C., and A.J. contributed to the conception and design of the study; J.M.R, conducted the data analysis; J.M.R wrote the manuscript; J.M.R produced the figures and data visualisations; J.M.R, R.C., and A.J. contributed to manuscript internal critical review process and revisions. All authors read and approved the submitted version.

## Data Accessibility Statement

All data and code used in this study are available on the *UK Data Service ReShare* at □; Data Collection □.

## Acknowledgements

We would like to thank Martin Breed, Chris Bedwell, Vicky Peace, Rachael White, and Kate Robinson for helping to pilot test the questionnaires and for providing feedback on the design.

## Supplementary Materials, Appendix A

### Online survey questions

– What is your age?
– What is your gender?
– What is/was your main occupation?
– What is your level of education?
– What country do you live in?
– What is your postal/zip code?
– How many times do you visit any natural environments (e.g., parks, woodlands, the beach) in a typical week?
– Approximately how long would you spend in any natural environment per visit?
– Select all of the organisms that you consider to be microbes (micro-organisms):
– Do you consider viruses to be:
– Do you consider all other microbes (micro-organisms) to be:
– From the list below, choose 3 words that you think best describe microbes:
– How much do you agree or disagree with the following: *Select one for each line*
– I feel very connected to all living things and the earth
– I always think about how my actions affect the environment
– My relationship to nature is an important part of who I am
– My connection to nature and the environment is a part of my spirituality
– My ideal holiday/vacation spot would be a remote, wilderness area
– I take notice of wildlife wherever I am

## References

Arleevskaya MI, Aminov R, Brooks WH, Manukyan G, Renaudineau Y. Shaping of Human Immune System and Metabolic Processes by Viruses and Microorganisms. Front Microbiol. 2020, 10, 816.

Balloux F, van Dorp L. Q&A: What are pathogens, and what have they done to and for us?. BMC Biol. 2017 Dec;15(1):1–6.

Birch, J.; Rishbeth, C.; and Payne, S.R. Nature doesn’t judge you–how urban nature supports young people’s mental health and wellbeing in a diverse UK city. Health & Place, 2020, p.102296.

Callaghan A, McCombe G, Harrold A, McMeel C, Mills G, Moore-Cherry N, Cullen W. The impact of green spaces on mental health in urban settings: a scoping review. J Mental Health. 2020 Apr 18:1–5.

Capaldi CA, Dopko RL, Zelenski JM. The relationship between nature connectedness and happiness: a meta-analysis. Front Psychol. 2014 Sep 8;5:976.

Capaldi CA, Passmore HA, Nisbet EK, Zelenski JM, Dopko RL. Flourishing in nature: A review of the benefits of connecting with nature and its application as a wellbeing intervention. Int J Wellbeing. 2015 Dec 17;5(4).

Cavicchioli R, Ripple WJ, Timmis KN, Azam F, Bakken LR, Baylis M, Behrenfeld MJ, Boetius A, Boyd PW, Classen AT, Crowther TW. Scientists’ warning to humanity: microorganisms and climate change. Nat Rev Microbiol. 2019 Sep;17(9):569–86.

Chawla L. Childhood nature connection and constructive hope: A review of research on connecting with nature and coping with environmental loss. People and Nature. 2020 Sep;2(3):619–42.

Cheung, H.; Mazerolle, L.;Possingham, H.P.;Tam, K.P.; and Biggs, D. A methodological guide for translating study instruments in cross‐cultural research: Adapting the ‘connectedness to nature’ scale into Chinese. Methods Ecol Evol, 2020, 11, pp.1379–1387.

Choe, E.Y.; Jorgensen, A.; and Sheffield, D. Does a natural environment enhance the effectiveness of Mindfulness-Based Stress Reduction (MBSR)? Examining the mental health and wellbeing, and nature connectedness benefits. Landscape and Urban Planning, 2020, 202, p.103886.

DeWeerdt, S. How baby’s first microbes could be crucial to future health. Nature. 555, s18–19.

Dannemiller KC, Mendell MJ, Macher JM, Kumagai K, Bradman A, Holland N, Harley K, Eskenazi B, Peccia J. Next‐generation DNA sequencing reveals that low fungal diversity in house dust is associated with childhood asthma development. Indoor air 2014 24, 236–47.

Flies EJ, Clarke LJ, Brook BW, Jones P. Urbanisation reduces the abundance and diversity of airborne microbes-but what does that mean for our health? A systematic review. Sci Tot Environ, 2020 Jun 22;738:140337-.

Guerra CA, Heintz-Buschart A, Sikorski J, Chatzinotas A, Guerrero-Ramírez N, Cesarz S, Beaumelle L, Rillig MC, Maestre FT, Delgado-Baquerizo M, Buscot F. Blind spots in global soil biodiversity and ecosystem function research. Nat Comm. 2020 Aug 3;11(1):1–3.

Haahtela T. A biodiversity hypothesis. Allergy. 2019 Aug;74(8):1445–56.

Jun S, Drall K, Matenchuk B, McLean C, Nielsen C, Obiakor CV, Van der Leek A, Kozyrskyj A. Sanitization of Early Life and Microbial Dysbiosis. Challenges. 2018 Dec;9(2):43.

Kettner, H.; Gandy, S.;Haijen, E.C.; and Carhart-Harris, R.L. From egoism to ecoism: Psychedelics increase nature relatedness in a state-mediated and context-dependent manner. Int J Environ Res Pub Health, 2019, 16, p.5147.

Li Q. Effect of forest bathing trips on human immune function. Environ Health Prevent Med. 2010 Jan 1;15(1):9–17.

Li M, Fang A, Yu X, Zhang K, He Z, Wang C, Peng Y, Xiao F, Yang T, Zhang W, Zheng X. Microbially-driven sulfur cycling microbial communities in different mangrove sediments. Chemosphere. 2020 Oct 13:128597.

Liddicoat C, Sydnor H, Cando-Dumancela C, Dresken R, Liu J, Gellie NJ, Mills JG, Young JM, Weyrich LS, Hutchinson MR, Weinstein P. Naturally-diverse airborne environmental microbial exposures modulate the gut microbiome and may provide anxiolytic benefits in mice. Sci Total Environ 2020 701, 134684.

Mayer FS, Frantz CM. The connectedness to nature scale: A measure of individuals’ feeling in community with nature. J Environ Psychol. 2004 Dec 1;24(4):503–15.

Nisbet EK, Zelenski JM. The NR-6: a new brief measure of nature relatedness. Front Psychol. 2013 Nov 1;4:813.

Nisbet EK, Zelenski JM, Grandpierre Z. Mindfulness in nature enhances connectedness and mood. Ecopsychology. 2019 Jun 1;11(2):81–91.

Obery A, Bangert A. Exploring the influence of nature relatedness and perceived science knowledge on proenvironmental behavior. Educat Sci. 2017 Mar;7(1):17.

Parks J, McCandless L, Dharma C, Brook J, Turvey SE, Mandhane P, Becker AB, Kozyrskyj AL, Azad MB, Moraes TJ, Lefebvre DL. Association of use of cleaning products with respiratory health in a Canadian birth cohort. CMAJ. 2020 Feb 18;192(7):E154–61.

Prescott SL. A Butterfly Flaps its Wings: Extinction of Biological Experience and the Origins of Allergy. Ann Allergy, Asthma Immunol. 2020 May 29.

Qadir MI. and Yameen IA. Questionnaire Based Study about Association between Blood Oxygen Level and Mysophobia. Biomed. J Sci Tech Res. 2019, 14:1–3

Renz H, Skevaki C. Early life microbial exposures and allergy risks: opportunities for prevention. Nat Rev Immunol. 2020 Sep 11:1–5.

Richardson, M., Hunt, A., Hinds, J., Bragg, R., Fido, D., Petronzi, D., Barbett, L., Clitherow, T. and White, M. A measure of nature connectedness for children and adults: Validation, performance, and insights. Sustainability, 2019, 11:3250.

Robinson JM, Cameron R. The Holobiont Blindspot: Relating Host-Microbiome Interactions to Cognitive Biases and the Concept of the “Umwelt”. Front Psychol. 2020;11.

Robinson JM, Breed MF. The Lovebug Effect: Is the human biophilic drive influenced by interactions between the host, the environment, and the microbiome?. Sci Tot Environ. 2020 Feb 28:137626.

Robinson JM, Cando-Dumancela C, Liddicoat C, Weinstein P, Cameron R, Breed MF. Vertical Stratification in Urban Green Space Aerobiomes. Environ Health Persp. 2020 Nov 25;128(11):117008.

Rook GA, Martinelli R, Brunet LR. Innate immune responses to mycobacteria and the downregulation of atopic responses. Curr Opin Allergy Clin Immunol 2003 3, 337–42.

Rook GA. Regulation of the immune system by biodiversity from the natural environment: an ecosystem service essential to health. Proc Nat Acad Sci. 2013 Nov 12;110(46):18360–7.

Roslund MI, Puhakka R, Grönroos M, Nurminen N, Oikarinen S, Gazali AM, Cinek O, Kramná L, Siter N, Vari HK, Soininen L. Biodiversity intervention enhances immune regulation and health-associated commensal microbiota among daycare children. Sci Adv. 2020 Oct 1;6(42):eaba2578.

Selway CA, Mills JG, Weinstein P, Skelly C, Yadav S, Lowe A, Breed MF, Weyrich LS. Transfer of environmental microbes to the skin and respiratory tract of humans after urban green space exposure. Environ Int. 2020 Dec 1;145:106084.

Smart Survey. Smart Survey online surveys. Available at: https://www.smartsurvey.co.uk/. Accessed 01-03-2020.

Stein MM, Hrusch CL, Gozdz J, Igartua C, Pivniouk V, Murray SE, Ledford JG, Marques dos Santos M, Anderson RL, Metwali N, Neilson JW. Innate immunity and asthma risk in Amish and Hutterite farm children. N Engl J Med. 2016 375, 411–21.

Timmis K, Cavicchioli R, Garcia JL, Nogales B, Chavarría M, Stein L, McGenity TJ, Webster N, Singh BK, Handelsman J, de Lorenzo V. The urgent need for microbiology literacy in society. Environ Microbiol. 2019 May;21(5):1513–28.

Trivedi P, Leach JE, Tringe SG, Sa T, Singh BK. Plant–microbiome interactions: From community assembly to plant health. Nat Rev Microbiol. 2020 Nov;18(11):607–21.

Whitburn J, Linklater W, Abrahamse W. Meta‐analysis of human connection to nature and proenvironmental behavior. Conservation Biology. 2020 Feb;34(1):180–93.

Yeager RA, Smith TR, Bhatnagar A. Green environments and cardiovascular health. Trends Cardio Med. 2020 May 1;30(4):241–6.

Yau, K.K.Y.; and Loke, A.Y. Effects of forest bathing on pre-hypertensive and hypertensive adults: a review of the literature. Environ Health Prev Med, 2020, 25, pp.1–17. 9.

Zemke DM, Neal J, Shoemaker S, Kirsch K. Hotel cleanliness: will guests pay for enhanced disinfection?. Int J Contemp Hospit Manag. 2015 May 11.

Zobell CE, Rittenberg SC. Microbiology by numbers. Nat Rev Microbiol. 2011; 9:628.

